# Optogenetic activation of striatal D1/D2 medium spiny neurons differentially engages downstream connected areas beyond the basal ganglia

**DOI:** 10.1101/2021.03.23.436576

**Authors:** Christina Grimm, Stefan Frässle, Céline Steger, Lukas von Ziegler, Oliver Sturman, Noam Shemesh, Johannes Bohacek, Klaas Enno Stephan, Daniel Razansky, Nicole Wenderoth, Valerio Zerbi

## Abstract

The basal ganglia (BG) are a group of subcortical nuclei responsible for motor control, motor learning and executive function. Central to BG function are striatal medium spiny neurons (MSNs) expressing D1 and D2 dopamine receptors. D1 and D2 MSNs are typically considered functional antagonists that facilitate voluntary movements and inhibit competing motor patterns, respectively. While their opposite role is well documented for certain sensorimotor loops of the BG-thalamocortical network, it is unclear whether MSNs maintain a uniform functional role across the striatum and which influence they exert on brain areas outside the BG. Here, we addressed these questions by combining optogenetic activation of D1 and D2 MSNs in the mouse ventrolateral caudoputamen (vl CPu) with whole-brain functional MRI (fMRI) recordings. Neuronal excitation of either cell population in the vl CPu evoked distinct activity patterns in key regions of the BG-thalamocortical network including the pallidum, thalamus and motor cortex. Importantly, we report that striatal D1 and D2 MSN stimulation differentially engaged cerebellar and prefrontal regions. We characterised these long-range interactions by computational modelling of effective connectivity and confirmed that changes in D1 / D2 output drive functional relationships between regions within and beyond the BG. These results suggest a more complex functional organization of MSNs across the striatum than previously anticipated and provide evidence for the existence of an interconnected fronto - BG - cerebellar network modulated by striatal D1 and D2 MSNs.

**Graphical Abstract:** 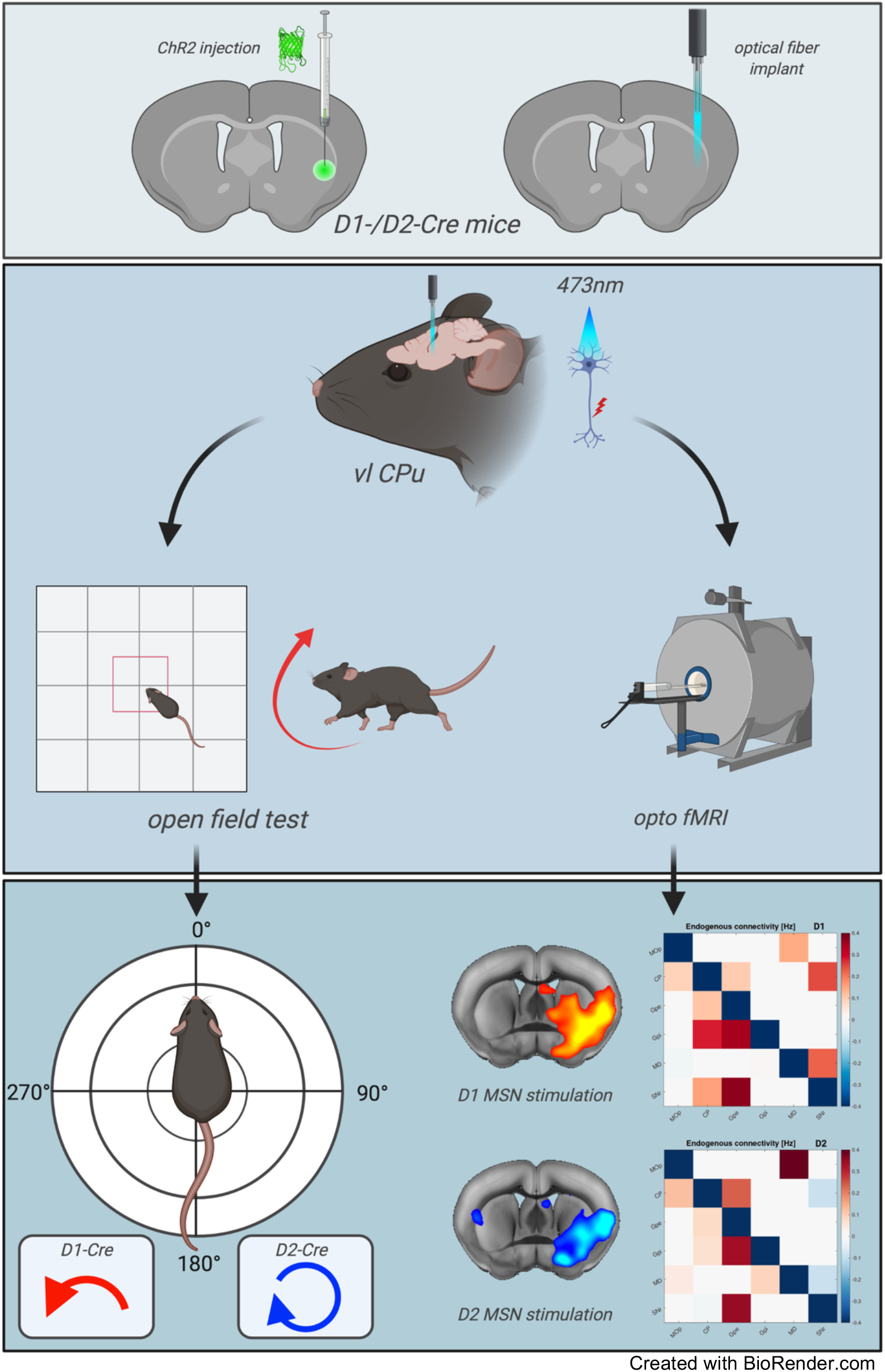

## Introduction

The basal ganglia (BG) integrate information from a wide array of cortical and thalamic inputs and take part in the regulation of motor, cognitive, and limbic functions [1]. An essential prerequisite for selection and execution of appropriate actions is the convergence of excitatory (glutamatergic) and modulatory (dopaminergic) afferents in the striatum. Striatal GABAergic medium spiny neurons (MSNs) expressing D1 and D2 dopamine receptors constitute the major recipients of those synaptic inputs. Traditionally, their role in the striatum is associated with motor function, conveyed via the direct and indirect pathway [1, 2]. D1 MSNs of the direct pathway directly innervate the GABAergic BG output nuclei - the substantia nigra pars reticulata (SNr) and internal globus pallidus (GPi) - which results in the disinhibition of thalamocortical circuits, allowing them to execute commands necessary for movement initiation. Conversely, D2 MSNs of the indirect pathway indirectly activate the SNr via the external globus pallidus (GPe) and the subthalamic nucleus (STN) resulting in the inhibition of thalamocortical circuits and a decrease in locomotor activity.

Although this canonical model is relevant for understanding BG-related disorders and therapeutic interventions, it assumes that MSNs maintain their functional role uniformly across the striatum. Contrary to this view, tracer studies in non-human primates and rodents have revealed a much more complex anatomical picture [3–6], with recent work counting as many as 29 anatomically distinct sub-regions in the mouse dorsomedial (dm) striatum (i.e. caudoputamen (CPu)) [7, 8]. Furthermore, anatomical connections in the form of collateral branches, recurrent networks and feedback loops suggest a causal influence of MSNs on brain regions beyond those considered by canonical BG models, such as cerebellum and prefrontal cortex [9–12]. So far, however, it remains elusive how D1 and D2 MSNs distinctly affect the function of the circuits inside and outside the BG and whether this depends on their anatomical location in the striatum. To date, only one study has investigated large-scale functional influences of cell type-specific activity in the mouse striatum and has provided direct evidence for the canonical model of BG motor function [13, 14]. However, this analysis was restricted to the dorsomedial sub-region of the CPu.

To gain further insight into how MSNs in other striatal sub-regions drive activity and functional interactions across the whole brain, we combined optogenetic stimulation of D1 and D2 MSNs in the mouse ventrolateral CPu (vl CPu) with whole-brain functional magnetic resonance imaging (fMRI) recordings and a model of effective connectivity (regression dynamic causal modelling, rDCM [15–17]). Our data show highly reproducible but different brain activation profiles during D1 and D2 MSN stimulation in the vl CPu. These results confirmed the causal role of D1 and D2 MSNs in BG function in line with the canonical model of the BG-thalamocortical network [13]. In addition, we observed a distinct involvement of the cerebellum and prefrontal regions depending on the stimulated MSNs. This information is useful for understanding the role and complexity of striatal D1 and D2 MSN output and for extending the canonical description of BG function to the whole brain.

## Methods

### Animals

All animal procedures were conducted in accordance with the Swiss Federal Ordinance on Animal Experimentation and approved by the Cantonal Veterinary Office of Zurich. Experiments were performed using 8-to 12-week-old male and female D1-Cre (n(male)=5; n(female)=6) and D2-Cre (n(male)=4; n(female)=4) mice. Animals were kept in standard housing on a 12h light/dark cycle. Food and water were provided *ad libitum*.

### Stereotaxic surgery

Viral vectors were obtained from the Viral Vector Facility (VVF) of the Neuroscience Center Zurich. Experimental D1- and D2-Cre mice were injected unilaterally with 300 nl of an AAV viral construct (AAV5-hEF1α-dlox-hChR2(H134R)_EYFP(rev)-dlox-WPRE-hGHp(A); titer: 6.3 × 10E12 vector genomes/ml). Control mice (D1-Cre) were injected with 300 nl of a fluorophore-only carrying AAV construct (AAV8-hSyn-dlox-EYFP(rev)-dlox-WPRE-hGHp(A); titer: 6.3 × 10E12 vector genomes/ml). Stereotaxic surgery was performed under anesthesia with a mixture of midazolam (5mg/ml; Sintetica, Switzerland), fentanyl (50mcg/ml; Actavis AG, Switzerland) and medetomidine (1mg/ml; Orion Pharma, Finland). Animals were placed into a stereotaxic apparatus (Neurostar, Germany) and their skulls exposed. Etching gel (DMG, Germany) was applied to the air-dried skull for 30 seconds and the bone surface was gently scratched using a metal scaler. After removal of bone debris with saline, bregma was located and the skull placement corrected for tilt and scaling. For virus delivery and optical fiber implantation in the ventrolateral striatum a small hole was drilled at AP −0.1, ML −2.6, DV 3.7 relative to bregma. Following virus delivery, an optical fiber bent to a 90-degree angle (400μm, NA=0.66; Doric Lenses, Canada) was placed at 0.2 mm above the targeted site. 90 degree bent optical fibers, optimized to fit the high SNR cryogenic MRI coil, were glued to the skull using a UV-curable dental composite (Permaplast, LH Flow; M+W Dental, Germany) and stitches were used as required. Throughout the surgical procedure, body temperature was kept at 35 °C using a heating pad (Harvard Apparatus, USA). Following implantation, an anesthesia antidote mixture (temgesic (0.3mg/ml; Reckitt Benckiser AG, Switzerland), annexate (0.1mg/ml; Swissmedic, Switzerland), antisedan (0.1mg/ml; Orion Pharma, Finland) was administered subcutaneously and mice were placed into a heating chamber to recover. All animals were monitored for 3 days after surgery and left to recover for 3 - 4 weeks before behavioural and scanning sessions.

### Open-field test in freely moving mice

Behavioural testing took place in a dimly lit chamber with sound insulating properties. Laser-light evoked changes in rotational behaviour of experimental D1- and D2-Cre mice [18, 19] were quantified during an open-field test. Animals were placed in a 40 × 40 cm arena and left to explore freely for 15 minutes. After this habituation period, mice were subjected to five cycles of 20s laser stimulation at 473 nm and 20 Hz (ON) with 5mW laser power followed by 40s of no laser stimulation (OFF). Motor behaviour was video recorded at all times.

### Opto-fMRI recording

Data were acquired in a 7T Bruker BioSpec scanner equipped with a Pharmascan magnet and a high SNR dedicated mouse brain cryogenic coil (Bruker BioSpin AG, Fällanden, Switzerland).

#### Animal preparation

For fMRI data acquisition, mice were anesthetized in a gas chamber for 4 minutes with 4% isoflurane in 1:4 O2 to air mixture. Animals were endotracheally intubated and the tail vein cannulated while being kept under anesthesia with 2% isoflurane. During preparation, animal temperature was kept at 35 °C using a heating pad (Harvard Apparatus, USA). Once intubated and cannulated, mice were head-fixed with earbars and connected to a small animal ventilator (CWE, Ardmore, USA) on an MRI-compatible support. Ventilation was set to 80 breaths per minute, with 1.8 ml/min flow with isoflurane at 2%. A bolus containing a mixture of medetomidine (0.05mg/kg) and pancuronium (0.25mg/kg) was delivered via the cannulated vein. Isoflurane was set to 1.5%. Continuous infusion of medetomidine (0.1mg/kg/h) and pancuronium (0.25mg/kg/h) started five minutes after the initial bolus injection. Isoflurane was reduced to 0.5%. A hot water-circulation bed kept the temperature of the animal constant throughout the entire measurement (36 °C). After collection of functional and anatomical fMRI scans continuous injection and isoflurane flow were stopped. Animals remained connected to the ventilator until independent breathing could be assured. For further recovery, they were transferred to a heating chamber.

#### fMRI sequences

For each experimental mouse, we acquired three different modalities of functional acquisition in two fMRI scanning sessions. Gradient-echo (GE) and Spin-echo (SE) sequences were acquired to capture BOLD contrast via T2* and T2 weighted signals. In addition, we implemented isotropic diffusion encoding (IDE) gradient waveforms to impart a diffusion weighting functional contrast as in [20, 21]. Control mice were scanned using the GE BOLD fMRI sequence only. For GE BOLD fMRI, an echo planar imaging sequence (EPI, repetition time TR = 1000 ms, total volumes = 480, slice thickness ST = 0.45 mm, in-plane spatial resolution RES = 0.22×0.2 mm^2^) was used to collect 2 to 4 datasets from each mouse. For SE sequences (including the IDE gradient waveforms implementation for diffusion weigthing), 4 datasets were acquired in each mouse within a single session. For diffusion-weighted fMRI (dfMRI), a SE EPI sequence was used (TR = 1000 ms, total volumes = 480, ST = 1.45 mm, RES = 0.23×0.23 mm^2^, b = 1500 s/mm^2^). To impart BOLD contrast, this sequence was used as is (i.e. b = 0 s/mm^2^), given it delivers T_2_ - weighted signals [22]. For both SE and dfMRI sequences, EPI slices where positioned according to anatomy, capturing the entire midbrain and part of the PFC (bregma +1.33).

Anatomical scans were acquired via a FLASH sequence with an in-plane resolution of 0.05 × 0.02 mm^2^, an echo time (TE) of 3.51 ms and a repetition time (TR) of 522 ms.

#### Optogenetic stimulation

Experimental and control mice were light stimulated via an optical fiber (Doric Lenses, Canada) connected to a custom-made DPSS laser (CNI laser, China). Following a baseline period of 170 seconds, trains of 473 nm laser pulses were delivered at 20 Hz and 5 mW for 20 seconds (ON) followed by 40 seconds of no laser light delivery (OFF) repeated over five minutes. The precise onset of laser pulses was controlled using the COSplay trigger device [23]. To mitigate the visual artifacts of ‘spilling’ blue laser light, an additional light source in the form of an LED lamp was placed on the cradle inside the scanner.

### Tissue collection and immunohistochemistry

Tissue collection for immunohistochemistry was performed at 90 min following a 3 min continuous 20 Hz optogenetic stimulation at 473 nm and 5 mW. Animals were deeply anesthetized with a ketamine/xylazine/acepromazine (100 mg/mL, 20 mg/mL, 10 mg/mL) mixture and intracardially perfused using 20 mL ice-cold PBS (pH 7.4). Brain tissue was fixated with an intracardial perfusion of ice-cold 4% PFA. The brain was dissected, post-fixated in 4% PFA at 4°C for 1 hour, rinsed with PBS and transferred to a sucrose solution (30% sucrose in PBS) to be stored at 4°C overnight. Once fully dehydrated, the brain was frozen in tissue mounting medium (Tissue-Tek O.C.T Compound, Sakura Finetek Europe B.V., Netherlands) and coronally sectioned into 40 μm thick slices using a cryostat (Leica CM3050 S, Leica Biosystems Nussloch GmbH). Brain sections were transferred to ice-cold PBS for further immunohistochemical processing. For immunohistochemistry, brain slices were submerged in a primary antibody solution of 0.2% Triton X-100 and 2% normal goat serum in PBS. Sections were incubated under continuous agitation at 4°C for 2 nights. After washing 3 times/10 minutes in PBS, sections were transferred to incubate in secondary antibody solution for 1 hour at room temperature. Brain sections were washed 3 times/10 minutes, mounted onto glass slides (Menzel-Gläser SUPERFROST PLUS, Thermo Scientific) and air-dried before cover-slipping with Dako fluorescence mounting medium (Agilent Technologies). Antibodies included rabbit anti-pre-pro-enkephalin (ppENK; 1:200, Neuromics Cat# RA14124 RRID:AB_2532106) with secondary antibody Alexa Fluor^®^ 546 goat anti-rabbit (A11035, Life Technologies, 1:300), rabbit anti-cFos (226 003, Synaptic Systems, 1:5000) with secondary antibody Alexa 546 goat anti-rabbit (A11035, Life Technologies, 1:300), rabbit anti-prodynorphin (PA5-96439, Thermo Fisher Scientific, 1:200) with secondary antibody Alexa Fluor^®^ 546 goat anti-rabbit (A11035, Life Technologies, 1:300), and Nissl stain (N21483, NeuroTrace 640/660 Nissl stain, Thermo Fisher Scientific, 1:300). Opsin and cFos expression were validated using a confocal laser-scanning microscope (CLSM 880, Carl Zeiss AG, Germany).

### Quantification and Statistical Analysis

#### Open field test in freely moving mice

Head angle tracking in freely moving, experimental mice was performed with DeepLabCut 2.0.7 [24] using 13 body points of interest and 4 landmark points of interest within the open field arena. The network was trained using 10-20 frames from randomly selected videos for 250’000 - 1’030’000 iterations. The data generated by DeepLabCut was processed using custom R scripts (available online at https://github.com/ETHZ-INS/DLCAnalyzer). Full details can be found here [25].

#### cFos expression

Confocal images were imported to ImageJ by Fiji [26] and neurons positive for ChR2-EYFP, cFos or both were counted manually.

#### GLM statistical mapping

Preprocessing and functional data analysis was carried out using FSL FEAT (version 5.92, www.fmrib.ox.ac.uk/fsl) and in-house Matlab scripts. Pre-statistical processing included the following steps: Pre-processing of the BOLD data included discarding the first ten measurements to ensure achieving steady-state excitation, high pass filtering (with a cut-off of 90s), motion correction using MCFLIRT, spatial smoothing using a Gaussian kernel of FWHM 4mm and interleaved slice timing correction. Registration was first carried out to the respective T1-weighted image and then to a custom standard space template, using FLIRT. Time series statistical analysis was carried out using FILM with local autocorrelation correction. In first-level analyses, contrasts for laser stimulation versus no laser stimulation were computed for each individual, including standard motion parameters as covariates. Fixed-effects analysis was performed to generate second-level contrasts at the group level (contrast 1: increase in BOLD signal, contrast 2: decrease in BOLD signal; initial cluster forming threshold Z > 3.1, cluster extent significance threshold of p = 0.05), using the first-level analysis results.

Second-level statistical cope (contrast of parameter estimate) images (contrast 1: increase in BOLD signal) were then used to estimate the mean (paired) difference to obtain group-contrast beta maps (D1>D2, D1<D2). For visualization purposes, averaged group and group contrast activation maps were normalized to a high-resolution Allen Brain Institute (ABI) anatomical atlas using a greedy transformation.

#### Dimensionality reduction

To visualize the dynamics of whole-brain BOLD changes evoked by D1 / D2 MSN activation, we projected our BOLD-fMRI data onto a low-dimensional space. Briefly, time-varying BOLD signals were averaged across conditions and converted to a 2D matrix of [voxel * time]. This matrix was then fed into an unsupervised manifold learning technique (Uniform Manifold Approximation and Projection, UMAP), which preserves the global data structure and local relationships with neighbours. Point-to-point distances in the UMAP plots were then used to interpret the continuity (or discontinuity) of the fMRI activity patterns and to identify similarities in temporal and spatial profiles between individual stimulations [27].

#### rDCM analysis

Anatomical regions of interest (ROIs) for the regression dynamic causal modeling (rDCM) analysis were selected using an in-house Matlab script. In short, local activation maxima and minima (connected components of voxels) in D1 and D2 group z-stat maps (experimental groups) were identified via an imposed upper and lower threshold. Once thresholded, pixel connectivity was set to a 6-connected neighbourhood and finally transformed into unique clusters of regional maxima/minima. From D1 and D2 group z-stat maps, a total of 30 clusters were identified and anatomically labelled. BOLD signal time series were then extracted as the average signal of all voxels within each cluster. Extracted time series entered effective (directed) connectivity analysis using rDCM.

For the rDCM analysis, we made use of the open-source rDCM toolbox, which is freely available as part of the TAPAS software package (www.translationalneuromodeling.org/tapas). In brief, rDCM is a novel variant of DCM for fMRI [28] that enables whole-brain effective connectivity analyses by reformulating the numerically costly estimation of coupling parameters in differential equations of a classical linear DCM in the time domain into an efficiently solvable Bayesian linear regression in the frequency domain [15, 16].

Here, we applied two different strategies: First, we restricted ourselves to key components of the canonical BG-thalamocortical network: (i) primary motor area (MOp), (ii) caudoputamen (CPu), (iii) external globus pallidus (GPe), (iv) internal globus pallidus (GPi), (v) mediodorsal nucleus of thalamus (MD), and (vi) substantia nigra (SNr). In a second step, we then assessed effective connectivity within a more extended brain-wide network that comprised 30 nodes, including cerebellar and prefrontal regions (among others).

For both analyses, we utilized the embedded sparsity constraints of rDCM to automatically prune connections without having to rely on *a priori* restrictions on network architecture [15]. To this end, we initially assumed fully (all-to-all) connected networks, where all brain regions were coupled via reciprocal connections. Starting from this fully connected network, model inversion then automatically pruned connections to yield sparse effective connectivity patterns. Notably, since exact a priori knowledge about the degree of sparseness of the networks was not available, we followed previously established procedures for obtaining an optimal estimate of the network’s sparseness 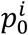 [15]. Specifically, for each mouse, we systematically varied 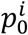 within a range of 0.1 to 0.9 in steps of 0.05 and performed model inversion for each 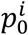 value. The optimal 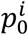 value was then determined for each mouse by selecting the setting that yielded the highest log model evidence (as approximated by the negative free energy).

Importantly, to assess the construct validity of our approach in the present context, we tested whether rDCM could correctly identify the region that received the optogenetic stimulation (i.e. vl CPu). This is possible (and meaningful) because the present setting represents a rare case where ‘ground truth’ (i.e. the region that received the optogenetic stimulation) is known in an empirical dataset. We challenged rDCM to faithfully recover this ground truth by creating a model space comprising 6 models for the BG network analysis and 30 models for the full network analysis, respectively. Models differed in the region that received the driving input (i.e. optogenetic stimulation). Specifically, model 1 hypothesized region 1 (MOp) to receive the driving input, model 2 hypothesized region 2 (vl CPu) to receive the driving input, and so on. We then utilized fixed-effects Bayesian model selection [29] to compare the competing models based on their model evidence.

For the winning model, we then inspected the posterior parameter estimates by performing fixed-effects Bayesian averaging to obtain group posterior distributions over model parameters. Significance of parameter estimates was then assessed in terms of their posterior probability exceeding a threshold of 0.95 [30].

## Results

### Optogenetic targeting of D1 and D2 MSNs in the ventrolateral striatum

For selective, light-driven activation of D1 and D2 MSNs, we stereotaxically injected a virus carrying a floxed, excitatory opsin in the right vl CPu of D1- and D2-Cre mice (**Figure 1A**) [19]. A 90-degree bent optical fiber was implanted above the targeted region to enable laser-light delivery. Histological staining confirmed that viral expression of the opsin ChR2 was located to either D1 or D2 MSNs co-expressing prodynorphin (DYN) and enkephalin (ENK), respectively (**Figure 1A**). Next, we sought to assess the behavioural relevance of light-driven increased D1 and D2 MSN activity in the open-field test (OFT). Previous reports of optogenetic MSN stimulation in the right dm CPu describe an increase in ipsiversive and contraversive rotations in D2- and D1-Cre mice, respectively [13, 31], which we aimed to replicate. After an acclimatisation period of 15 minutes, 5 cycles of 20 sec, 20 Hz laser pulses at 5 mW (ON) were delivered to the targeted MSN sub-population interleaved by 40 sec periods (OFF) without laser stimulation. Our results show a significant difference in head angle rotations specific to D1 vs D2 MSN stimulation (p = 0.00024, linear mixed effects model with *post hoc* test; **Figure 1C**). In D2-Cre mice, significant changes in ipsiversive median head angle position (p = 0.003, linear mixed effects model with *post hoc* test; **Figure 1C**) were accompanied by a significant increase in ipsiversive rotations (p = 0.012, two-tailed paired t-test; **Figure 1B**). In D1-Cre mice, laser stimulation evoked dystonic, contraversive changes in median head angle (p = 0.06, linear mixed effects model with *post hoc* test; **Figure 1C**) which coincided with a significant drop in average speed (p = 0.046, linear mixed effects model with *post hoc* test; **Supplementary Figure 1B**). These data indicate that optogenetic stimulation of D1 and D2 MSNs was sufficient to elicit changes at the behavioural level and show that movement patterns evoked by increased activity of D1 MSNs in the vl CPu differ from those observed when stimulating in the dm CPu.

**Figure 1.**
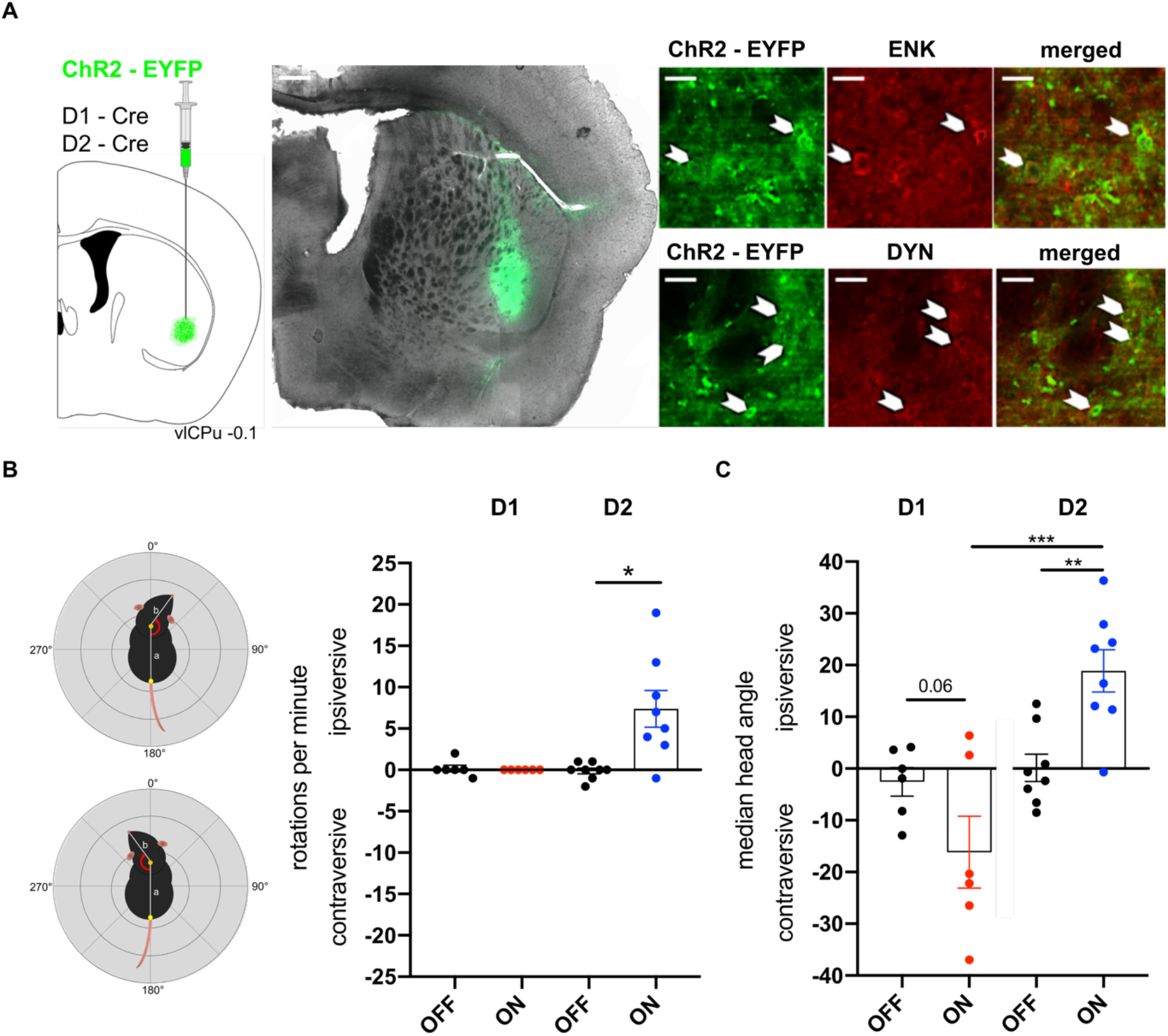
Optogenetically evoked D1/D2 MSN activity in the vl CPu. **A** Histological verification of D1 and D2 MSN targeting in the vl CPu. Arrowheads indicate cells co-expressing ChR2-EYFP and the stained antibody DYN and ENK in D1 and D2-Cre mice, respectively. **B** Optogenetically evoked motor behaviour measured as number of rotations per minute in the open-field test. In D1-Cre mice, rotational behaviour did not increase during laser stimulation (n = 6 animals; p = 0.695, two-tailed paired t-test). In D2-Cre mice, the number of ipsiversive rotations per minute significantly increased during laser stimulation (n = 8 animals, *p = 0.012, two-tailed paired t-test). **C** Optogenetically evoked moto-behavioural effects on median head angle position in the open-field test. Optogenetic stimulation evoked highly significant changes in median head angle in D1 vs D2-Cre mice (***p = 0.00024, linear mixed effects model with *post hoc* test). Specifically, laser stimulation in D2-Cre mice evoked significant changes in the head angle position in an ipsiversive manner (**p = 0.003, linear mixed effects model with *post hoc* test), while D1-Cre mice changed their head angle in an contraversive manner (p = 0.06, linear mixed effects model with *post hoc* test). DYN, dynorphin; ENK, encephalin; **p < 0.01, ***p < 0.001. Scale bars, 50μm, 500μm.

Following behavioural characterization, we proceeded to validate ChR2 activity on a molecular level using the neural activity marker cFos. Lightly anesthetised D1- and D2-Cre mice were stimulated for 3 min with continuous 20 Hz laser pulses at 473 nm and 5 mW. After 90 min, their brains were collected to immunohistochemically label cFos positive MSNs. In both D1- and D2-Cre mice, cFos expression co-localized with the majority of ChR2-EYFP positive cells (**Supplementary Figure 1D-G**) suggesting that our stimulation protocol was sufficient to elicit changes at the molecular level.

### Striatal D1 and D2 MSNs stimulation differentially engages downstream connected brain regions in and beyond the basal ganglia

To gain insight into downstream influences of either MSN sub-population in the vl CPu, we measured whole-brain responses induced by optogenetic stimulation of D1/D2 MSNs using BOLD fMRI. Correct fiber placement to the vl CPu was verified for each mouse via T1-weighted anatomical scans (**Supplementary Figure 2A**). For the acquisition of functional scans, 5 cycles of 20 sec blue-light laser pulse trains at 20 Hz (ON) followed by a 40 sec post-stimulation period (OFF) were applied to evoke robust brain-wide activation patterns (**Figure 2A**). D1-Cre mice expressing no opsin but implanted with an optical fiber in the right vl CPu were used as controls. Using a GLM approach, we identified the brain areas that were significantly modulated by the laser stimulation with voxel resolution (200um isotropic; **Figure 2B**) and extracted their mean fMRI time-series (**Figure 2C** and **Supplementary Figure 3**, see Methods for the definition of local maxima/minima). Group activation maps showed brain-wide BOLD activity patterns evoked by D1 and D2 MSN stimulation. We observed strong and nearly identical BOLD signal increases in the target region (right vl CPu) (**Figure 2C upper left panel**), suggesting that we successfully matched stimulation strength in both conditions. Importantly, no BOLD changes were recorded in the vl CPu of control mice (**Supplementary Figure 2B**). Key regions connected to the BG like the thalamus (TH) and primary motor cortex (MOp) exhibited BOLD activation patterns in line with the canonical description of the direct pathway (**Figure 2B** and **Supplementary Figure 3A**). Contrary to the predictions of the canonical model, stimulating D1 MSNs evoked positive BOLD signal changes in the STN and GPi (**Figure 2B**). Further discrepancies with the canonical model were recorded in the GPe, where positive BOLD signal changes opposed its presumed inhibition during D2 MSN stimulation (**Supplementary Figure 3A**). Beyond the BG, D1 and D2 MSN stimulated mice showed comparable BOLD profiles in the ipsilateral agranular insula (AI), the anterior amygdalar area (AAA) and the temporal association area (TEa) (**Supplementary Figures 2C** and **3B**), while other extra-BG regions exhibited differences in their BOLD activity (**Figure 2B-C**): D2 MSN activation resulted in positive BOLD responses in the anterior cingulate (ACAv) and infralimbic (IL) area of the mPFC, the dentate gyrus (DG) and the cerebellar Crus I region. Instead, the midbrain reticular nucleus (MRN) and motor-related superior colliculus (SCm) as well as the cerebellar simple lobule (SIM) showed increased BOLD activity upon D1 MSN stimulation (**Figure 2B-C**). These results provide evidence of D1 and D2 MSN activity-dependent, canonical and non-canonical modulation of distinct brain regions within and beyond the BG.

**Figure 2.**
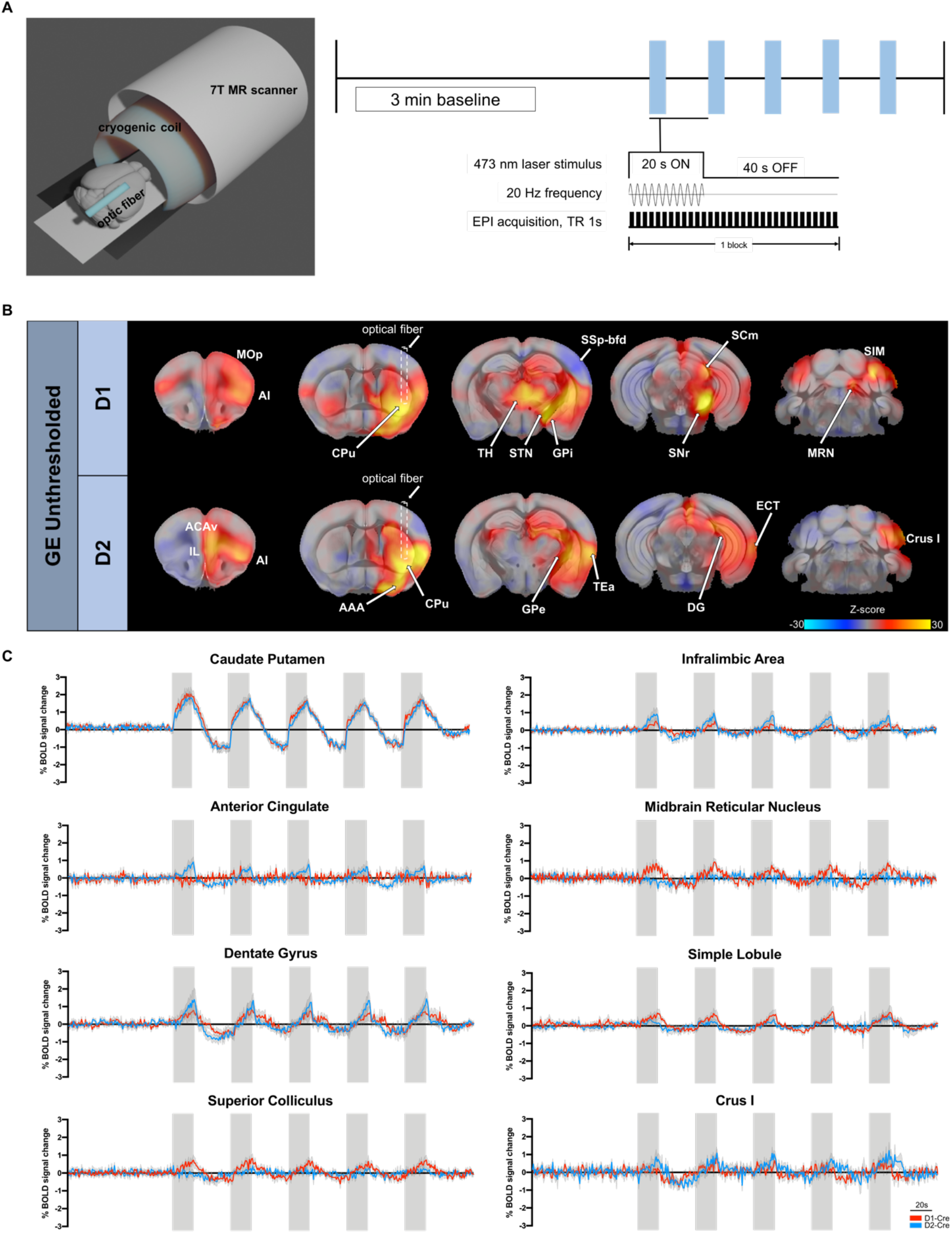
Opto-fMRI reveals causal brain-wide influences of D1 and D2 MSN activity. **A** Schematic opto-fMRI set-up and stimulation protocol. **B** Unthresholded GLM z-stat activation maps of D1 and D2 MSN stimulation. **C** Mean time-series of D1-/D2-Cre mice extracted from selected regions of interest based on GE BOLD local activation maxima and minima z-stat maps. D1 MSN time-series depicted in red, D2 MSN time-series depicted in blue. Laser stimulation blocks indicated in grey. TR, repetition time; EPI, echo planar imaging; AAA, anterior amygdalar area; AI, agranular insula; ACAv, anterior cingulate; CPu, caudate putamen; TH, thalamus; SNr, substantia nigra; SSp-bfd, primary somatosensory cortex, barrel field; SCm, superior colliculus, motor related; STN, subthalamic nucleus; TEa, temporal association area; ECT, ectorhinal area; IL, infralimbic area; GPe, external globus pallidus; DG, dentate gyrus; MOp, primary motor cortex; MRN, midbrain reticular nucleus; SIM, simple lobule.

### Striatal D1 and D2 MSN stimulation differentially alters dynamics of functional brain responses

After having characterized the spatial extent of D1/D2 MSN activation, we focused on further elucidating the dynamics of these functional brain responses. One way to do this is to use dimensionality reduction techniques [32]. These approaches can uncover latent functional patterns in complex datasets by distilling high-dimensional brain activity patterns into a small number of low-dimensional spatiotemporal components [33]. After concatenating the datasets, each repetition time block (1 second) was projected onto a low-dimensional space using an unsupervised manifold learning technique (Uniform Manifold Approximation and Projection, UMAP) [27]. The point-to-point trajectories in the UMAP plots were then used to interpret the dynamics of the D1/D2 MSN stimulations and to evaluate the reproducibility of the activation patterns across the five stimulation blocks. D1 and D2 MSN datasets showed a response to the optogenetic stimulation in three and four phases, respectively (**Figure 3A-B**). Importantly, for D1 and D2 stimulation, the observed patterns formed distinct trajectories in the low-dimensional manifold, indicating that their activation engages different brain areas (**Figure 3C-D, Supplementary Videos 1-2**). Both stimulations evoked an initial rapid response to the delivered laser stimulus lasting 3-4 seconds (phase 1) followed by a sustained response for the remainder of the stimulation (15-16 seconds, phase 2; **Figure 3 A-B**). However, while in D1 MSN datasets BOLD activity gradually returned to baseline levels once laser stimulation was terminated (39-40 seconds, phase 3; **Figure 3A)**, D2 MSN datasets showed a different dynamic response driven by a rapid drop in BOLD activity of prefrontal and cerebellar regions (1-2 seconds, phase 3; **Figure 3B** and **Supplementary Video 2**). Thereafter, D2 MSN datasets gradually returned to baseline levels (38-39 seconds, phase 4; **Figure 3B**). This indicates not only the engagement of different brain regions, but also differences in brain-wide dynamics during D1 and D2 MSN stimulation.

**Figure 3.**
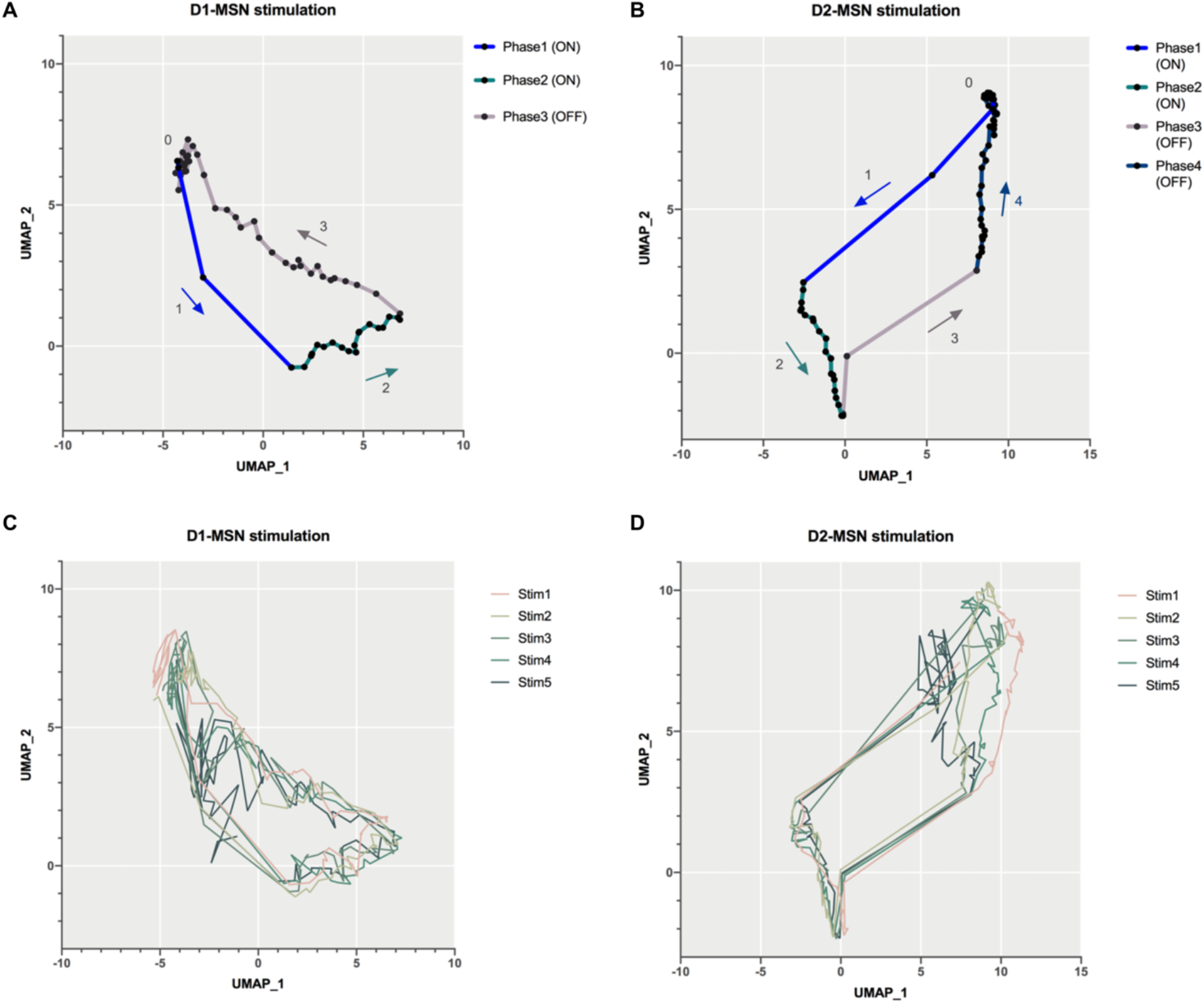
Functional brain-wide responses dynamically change upon D1 and D2 MSN stimulation. **A** Low-dimensional representation of averaged stimulation blocks in D1 MSN fMRI datasets. **B** Low-dimensional representation of averaged stimulation blocks in D2 MSN fMRI datasets. **C** Low-dimensional representation of single stimulation blocks in D1 MSN fMRI datasets. **D** Low-dimensional representation of single stimulation blocks in D2 MSN fMRI datasets. UMAP, Uniform Manifold Approximation and Projection. **Supplementary Video 1** Functional brain-wide responses dynamically change during D1 MSN stimulation. Demeaned and thresholded D1 MSN GE BOLD datasets visualized in red colour code. **Supplementary Video 2** Functional brain-wide responses dynamically change during D2 MSN stimulation. Demeaned and thresholded D2 MSN GE BOLD datasets visualized in blue colour code.

### Different fMRI contrasts capture robust D1 and D2 MSN brain-wide activation patterns

In addition to the classical GE-BOLD, new fMRI sequences allow to adjust the sensitivity of scans to their contrast types, which in turn influences functional activation patterns. Here, we rescanned experimental mice using two additional fMRI sequences with different intrinsic contrasts; namely, a spin-echo (SE) which enhances BOLD contrast from microcapillaries [34–36] and a diffusion-weighted functional scan (dfMRI) that is sensitive to changes in the intra- and extravascular diffusion of water molecules upon neural activation [22, 37]. SE and dfMRI sequences alike captured D1 and D2 MSN activity-dependent fMRI signal changes in key regions within the BG (**Supplementary Figure 4**).

We next aimed to obtain a more precise anatomical delineation of the difference between D1/D2 MSN activation profiles. To do that, we compared D1 and D2 datasets in a consensus analysis across all three different functional contrasts. In short, contrast estimates based on GE, SE and dfMRI acquisitions were pooled at the group level to compute a between-group statistical map, harboring group-specific activation across all three functional contrasts (**Figure 4A**). By comparing the brain-wide influences of each MSN sub-population, we observed that D1 MSNs drive stronger activity in the MOp, the MRN, the ipsi- and contralateral SIM as well as in the ipsi- and contralateral SSp-m, while D2 MSNs more potently activated the ipsilateral Crus I of the cerebellum, the majority of the ipsilateral DG, the ACAv and the IL area of the mPFC (**Figure 4B**). Additionally, D2 MSN stimulation elicited greater engagement of the ipsilateral SSp, including the upper and lower limb, trunk and barrel field area compared to D1 MSN activity. These findings further corroborate the different roles of D1 and D2 MSNs in driving brain-wide dynamics in and outside the BG.

**Figure 4.**
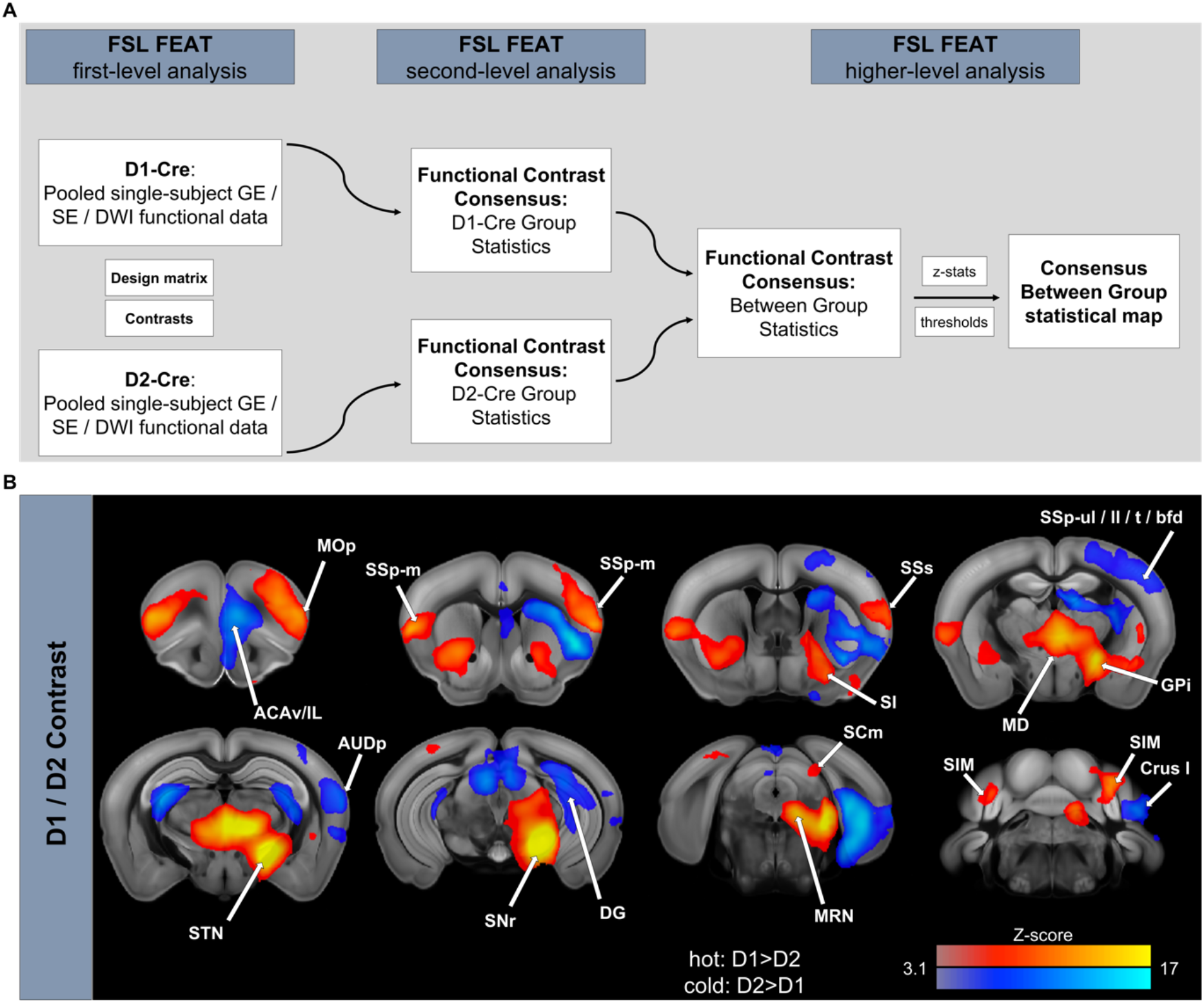
D1 and D2 MSNs differentially activate brain regions beyond the BG. **A** D1 and D2-Cre functional data analysis to create a GE / SE / dfMRI consensus between-group z-stat activation map using FSL FEAT. **B** GE / SE / dfMRI consensus z-stat activation map (cluster corrected; p<0.05). ACAv, anterior cingulate; IL, infralimbic area; MOp, primary motor cortex; SSp-m, primary somatosensory cortex, mouth area; SSp-ul, primary somatosensory cortex, upper limb area; SSp-ll, primary somatosensory cortex, lower limb area; SSp-t, primary somatosensory cortex, trunk area; SSp-bfd, primary somatosensory cortex, barrel field area; SI, substantia inominata; SSs, supplementary somatosensory cortex; GPi, internal globus pallidus; MD, mediodorsal nucleus of thalamus; STN, subthalamic nucleus; AUDp, primary auditory area; DG, dentate gyrus; SNr, substantia nigra; SCm, superior colliculus, motor related; MRN, midbrain reticular nucleus; SIM, simple lobule. Color code hot (red to bright yellow): D1>D2; color code cold (blue to turquoise): D2>D1.

### Regression dynamic causal modeling reveals distinct network connectivity participation upon D1 and D2 MSN stimulation

Traditionally, neuronal circuit models are based on the fragmented analysis of anatomical connections and individually acquired electrophysiological parameters. Novel techniques, however, strive to achieve a more integrative perspective on functionally connected neural networks by uniting whole-brain functional recordings, neurostimulation and computational modeling [14]. To infer upon the directed interactions amongst brain regions (i.e. effective connectivity) within and beyond the BG-thalamocortical network, we applied regression dynamic causal modeling (rDCM) to our opto-fMRI data.

First, we assessed effective connectivity among six key components of the canonical BG-thalamocortical network (MOp, CPu, GPe, GPi, MD, and SNr). Reassuringly, for both D1 and D2 MSN stimulation, rDCM correctly identified the vl CPu as the input region. More specifically, using fixed-effects Bayesian model selection (BMS) to compare all possible hypotheses of which region received the optogenetic stimulation yielded decisive evidence for the correct model with vl CPu as input region (posterior model probability of 1 for both D1 / D2 MSNs).

Inspecting the group-level posterior parameter estimates of the winning model, we found the effective connectivity patterns to be consistent with previous studies on the functional integration in the BG-thalamocortical network [14]. In brief, during D1 MSN stimulation (**Figure 5A**), we observed strong efferent connections from vl CPu to GPi and SNr, but not to GPe. Furthermore, strong connections were observed from SNr to GPe and GPi. Finally, the BG-output nucleus SNr sent strong projections to MD, which in turn sent information to the motor-output region MOp. Conversely, during D2 MSN stimulation, functional integration was overall weaker than during D1 MSN stimulation and displayed a distinct pattern (**Figure 5B**). When directly comparing D1 and D2, we found connections from vl CPu to GPi and from vl CPu to SNr – the defining projections of the direct pathway – to be significantly greater during D1 as compared to D2 MSN stimulation (**Figure 5C**). Conversely, the connection from vl CPu to GPe, a key region of the indirect pathway, was significantly increased during D2 as compared to D1 MSN stimulation. Furthermore, functional integration between GPi and SNr, as well as from the BG-output nucleus SNr to GPe and MD was greater during D1 as compared to D2 MSN stimulation. In addition to the aforementioned differences, other connections showed a significant differential effect of the optogenetic stimulation (**Figure 5C**).

**Figure 5.**
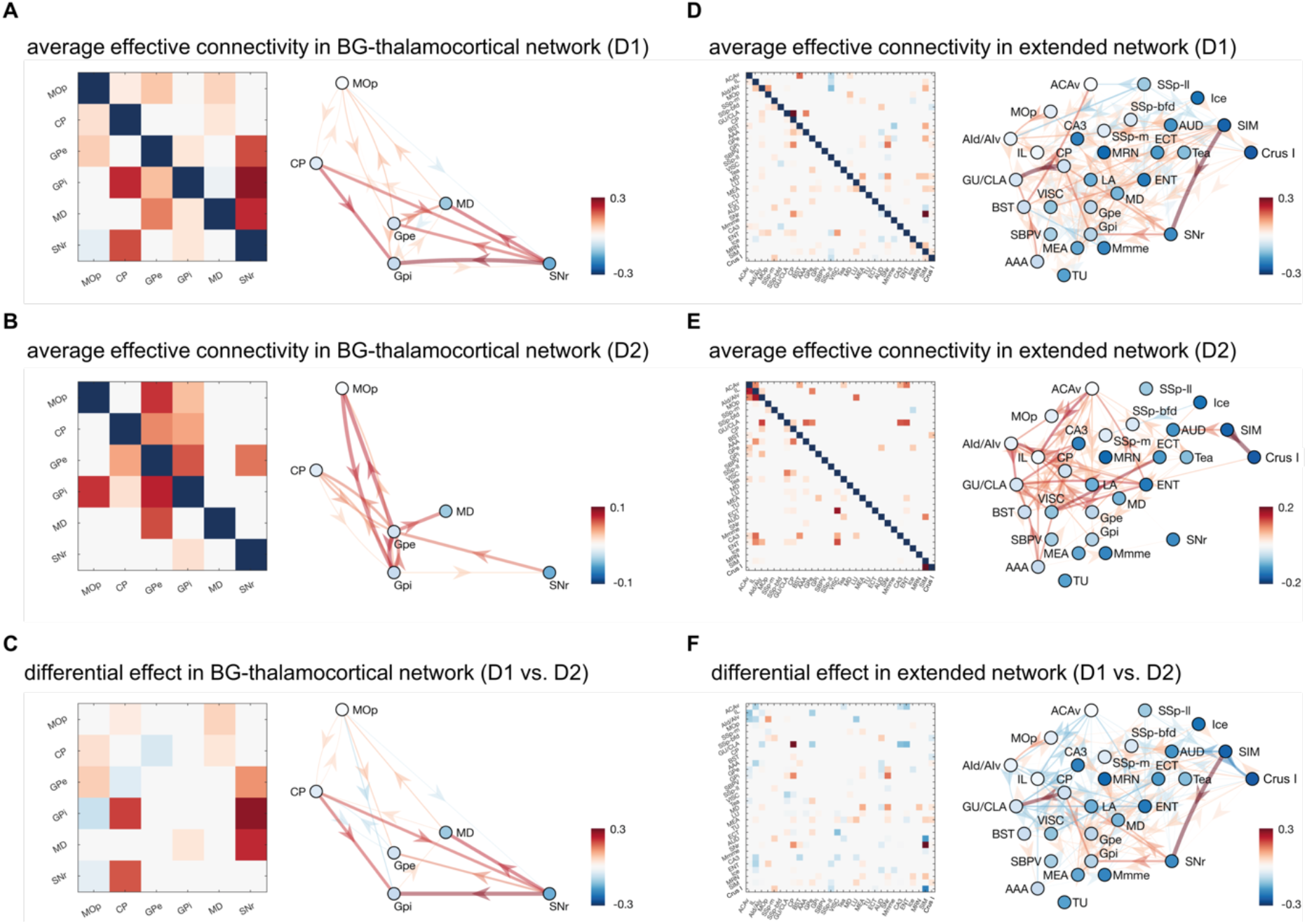
Effective (directed) connectivity during D1 or D2 MSN stimulation as inferred using regression DCM (rDCM). **A** Average connectivity pattern in the BG-thalamocortical network during D1 MSN stimulation, and **B** during D2 MSN stimulation (Pp > 0.95). **C** Differential effect of the optogenetic stimulation (D1-D2) in the BG-thalamocortical network (Pp > 0.95), where red colors indicate stronger connections during D1 stimulation and blue colors indicate stronger connections during D2 stimulation. **D** Average connectivity pattern in the extended network during D1 MSN stimulation, and **E** during D2 MSN stimulation (Pp > 0.95). **F** Differential effect of the optogenetic stimulation (D1-D2) in the extended network (Pp > 0.95), where red colors indicate stronger connections during D1 stimulation and blue colors indicate stronger connections during D2 stimulation. Note, color maps of each subplot have been adjusted to the respective maximal absolute connection strength and may thus differ between subplots. AAA, anterior amygdalar area; ACAv, anterior cingulate; PrL, prelimbic area; AIv, ventral agranular insula; AUD, auditory area; BST, Bed nuclei of the stria terminalis; CA3, Ammon’s horn field 3; CPu, caudate putamen; ECT, ectorhinal area; ENT, entorhinal area; GPe, external globus pallidus; GPi, internal globus pallidus; GU/CLA, gustatory area/claustrum; Ice, inferior colliculus; IL, infralimbic area; LA, lateral amygdala nucleus; MEA, medial amygdala nucleus; MD, mediodorsal nucleus of thalamus; Mmme, medial part of the medial mammillary nucleus; MOp, primary motor cortex; MRN, midbrain reticular nucleus; SIM, simple lobule; SNr, substantia nigra pars reticulata; SBPV, subparaventricular zone; SSp-bfd, primary somatosensory cortex, barrel field; SSp-ll, primary somatosensory cortex, lower limb area; SSp-m, primary somatosensory cortex, mouth area; TEa, temporal association area; TU, tuberal nucleus; VISC, visceral area.

In a next step, we investigated functional integration beyond the BG in an extended brain-wide network, including all regions in the mouse brain that were jointly activated or deactivated by the D1 and D2 MSN stimulation. Again, we first verified that rDCM correctly identified vl CPu as the target region that received the optogenetic stimulation for both D1 (FFX BMS: posterior model probability of 0.88) and D2 (FFX BMS: posterior model probability of 1).

Inspecting the group-level posterior parameter estimates of the winning model, we initially verified that, in this extended (whole-brain) analysis, the effective connectivity patterns in the BG-thalamocortical network were consistent with those obtained from the previous, more local analysis of connectivity. In brief, during D1 MSN stimulation (**Figure 5D**), we again observed strong engagement of the defining projections of the direct pathway, namely, the efferent connections from vl CPu to GPi and SNr (**Supplementary Figure 5A**), as well as strong connections from the BG-output nucleus SNr to MD (**Supplementary Figure 5B**). Going beyond the canonical BG-thalamocortical network, we observed strong projections from cerebellar regions, in particular SIM, to various components of the BG-thalamocortical network, in particular, SNr but also GPe and MD (**Supplementary Figure 5C**). Finally, we found strong efferent connections from vl CPu to GU/CLA (**Supplementary Figure 5A**). During D2 MSN stimulation (**Figure 5E**), the effective connectivity patterns again differed substantially from the D1 MSN stimulation. Specifically, as expected, we found no engagement of the direct pathway (**Supplementary Figure 5D**). Instead, we observed a strong involvement of prefrontal regions (e.g. AId/AIv, IL) as well as cingulate regions (e.g. ACAv) (**Supplementary Figure 5E**). Furthermore, we observed involvement of cerebellar structures (where the strongest projection was between two cerebellar regions, from SIM to Crus I) (**Supplementary Figure 5F**). Explicitly testing for difference between D1 and D2 revealed a wide range of differences both within the canonical BG-thalamocortical network and beyond, in prefrontal and cerebellar areas (**Figure 5F**).

In summary, rDCM provides further evidence that D1 and D2 MSNs drive activity in areas beyond the basal ganglia, involving cerebellar and prefrontal structures in a differential manner.

## Discussion

Traditionally, motor function of the BG is described as a feed-forward model, wherein D1 MSNs of the direct pathway disinhibit thalamo-cortical circuits necessary for movement initiation and D2 MSNs of the indirect pathway act opposingly via an increase of the BG’s inhibitory control over thalamus [1]. In line with this view, we found that activation of D1 but not D2 MSNs in the vl CPu elicited positive BOLD responses in key structures of the direct pathway including the TH and motor cortex. These results were further corroborated by effective connectivity analysis using regression DCM (rDCM), which highlighted the importance of the direct pathway during D1 but not D2 MSN stimulation. Using different fMRI contrast acquisitions, we were able to attenuate signal biases from putatively vascular sources and thereby provide a solid consensus on the organization of brain-wide activation patterns driven by either MSN population. Importantly, we observed different but not necessarily opposing effects of D1 and D2 MSN stimulation within the BG. Beyond the BG, D1 and D2 MSN stimulation differentially involved medial prefrontal regions as well as ipsi- and contralateral cerebellar lobules, which rDCM analysis also faithfully recapitulated. Our findings suggest a more complex functional organization of MSNs across the striatum than previously anticipated and provide evidence for the existence of an interconnected fronto - BG - cerebellar network modulated by striatal MSNs.

The behavioural output of striatal D1 and D2 MSNs is closely linked to motor initiation and suppression, respectively [38]. Rodent behavioural studies that stimulated D1 and D2 MSNs in the dm CPu have repeatedly supported their opposing roles [13, 31, 38, 39]. When tested in the OFT, unilateral stimulation of D2 MSNs in the vl CPu resulted in an increase of ipsiversive full body rotations; a behaviour that is identical to the one elicited during dm CPu D2 MSN stimulation [13, 31]. Instead, D1 MSN stimulation produced contraversive, dystonia-like head and neck movements. This differs from the full body contraversive, rotational phenotype reported in D1-Cre mice when stimulated in the dm CPu [13, 31]. In the dorsal CPu, electrophysiological studies have shown that D1 MSNs are less excitable than D2 MSNs and that concurrent activation of both MSN types is neccessary for contraversive movements [39, 40]. However, whether this difference between D1 and D2 MSNs is present in other sub-regions like the vl CPu and how this would affect motor behaviour is yet to be investigated. Overall, our data suggest a role of D1 MSNs on motor behavior that is heterogeneous and dependent on their anatomical location in the CPu, while for D2 MSNs, the exact location in the CPu does not seem to affect motor output.

On a more speculative note, the repetitive nature of head and neck movements in D1-Cre mice strongly resembled the clinical presentation of cervical dystonia, a syndrome of sustained or intermittent muscle contractions causing sideway twisting of the head [41]. Although relatively little is known about BG neuronal activity in dystonia, a recent study reported increased activity of the direct pathway in patients suffering from dystonic, involuntary movements: Simonyan et al. [42–44] found increased D1 receptor availability in the striatum of dystonic patients, which was paralleled by abnormally decreased D2 dopaminergic function via the indirect BG pathway. Ultimately, this imbalance between D1/D2 within the BG pathways could give rise to dystonic movement patterns [45]. Our findings are in line with these human findings and suggest that optogenetic stimulation of D1 MSNs in the vl CPu could induce such an imbalance in the BG and evoke dystonia-like movements also in mice. This assumption warrants a more thorough behavioural characterization of striatal D1 MSN hyperactivity and its potential role in dystonia.

The canonical model of BG function proposes opposing effects of D1 and D2 MSNs on BG-thalamocortical activity [1, 2]. Using opto-fMRI, we measured the cell type and region-specific influences of D1 and D2 MSNs and tested the validity of this model within the BG circuit. We further mapped the effective (directed) connectivity amongst key BG regions using rDCM [15, 16] to infer how either MSN sub-population contributes to reshaping the functional architecture of the BG-thalamocortical network. Importantly, effective connectivity among key regions of the direct pathway was significantly increased during D1 MSN stimulation while defining projections of the indirect pathway were significantly greater during D2 MSN stimulation, suggesting that their net output remains closely linked to ‘go’ / ‘no-go’ motor commands, respectively. Nevertheless, we found that BOLD dynamics could not support strictly antagonistic influences of vl CPu D1 and D2 MSNs within the BG-thalamocortical network. This included the TH and motor cortex. Here, D1 MSN stimulation elicited significant BOLD signal increases, but no changes were recorded during D2 MSN stimulation, which contradicted their presumed inhibition, i.e. de-activation during D2 MSN activity [1, 2]. Further, we found no opposing effects of D1 and D2 MSNs on BOLD activity in GPi. Here, positive BOLD signal changes were evoked only during D2 not D1 MSN stimulation. BOLD signals of the SNr and STN did not fit their predicted opposing profiles during D1 and D2 MSN stimulation either. These findings stand in contrast to previous opto-fMRI data by Lee et al. [13]. Targeting the dm CPu, they showed positive BG BOLD responses during D1 MSN stimulation, while negative BG BOLD responses were evoked during D2 MSN stimulation. Notably, while these responses did not strictly follow predictions of the canonical model either (see [13] for an extensive discussion thereof), they reflected the proposed antagonistic effects of D1 and D2 MSNs. Instead, our results indicate that during prolonged stimulation in the vl CPu, D1 and D2 MSNs do not exert opposing control but differentially engage activity in BG nuclei. This highlights the importance of considering the complex structural and functional architecture of the striatum when addressing the causal role of D1 and D2 MSNs within the BG.

The BG are traditionally modeled as an integrative structure which send and receive projections to each other and across a vast network of brain areas. Hence, we expected functional influences of striatal D1 and D2 MSNs to extend beyond the BG. In line with this view, we showed that D1 or D2 MSN activation led to changes in BOLD activity in the mPFC and cerebellar cortices. For example, D2 MSN excitation was followed by strong increases in BOLD signal in the ACAv and IL area of the mPFC, suggesting a causal influence of striatal D2 MSNs at this level. D2 MSN activity might participate in modulating the mPFC’s executive control of actions, especially those related to action inhibition [46, 47]. The mPFC has implications in a wide range of psychiatric diseases where executive control of action is compromised, including obsessive-compulsive disorder, addiction and anxiety [48, 49]. Importantly, separate lines of evidence indicate that the rodent mPFC-CPu circuit is relevant to anxiety and avoidance behaviour: using optogenetics and electrophysiology, Friedman and colleagues [50] demonstrated that ACAv to dm CPu projections are selectively active during an approach-avoidance T-maze task and that their excitation increases avoidance behaviour. Increased activity of mPFC-dmCPu projection neurons has also been reported during open arm exploration of an anxiogenic elevated zero maze (EZM) [51]. In fact, stimulation of D2 MSNs increased avoidance in the EZM and further heightened avoidance of open areas during an open field test [51, 52], suggesting that striatal D2 MSNs are critically involved in the control of avoidance behaviours [50–52]. Our results support a causal role of D2 MSNs in mPFC modulation and indicate that their functional influence is not limited to motor output but also involves higher cognitive processes and executive control.

We also found differential effects of D1 and D2 MSN stimulation in the cerebellum. D1 MSNs evoked BOLD activity in the bilateral SIM while D2 MSNs engaged the Crus I area. Along those lines, rDCM revealed that areas of the BG-thalamocortical network, in particular, SNr, GPe and MD, receive afferent inputs by SIM during D1 MSN stimulation. In contrast, intra-cerebellar projections from SIM to Crus I dominated during D2 MSN stimulation. Importantly, the cerebellum not only receives input and sends output to the prefrontal cortex [53, 54] but has also been shown to communicate with the BG via cerebellar motor and non-motor output nuclei. These nuclei project di-synaptically via the intralaminar thalamic nuclei to the striatum [10, 11] and ultimately give rise to an interconnected fronto - BG - cerebellar network. This network shows topographic organization: cognitive, limbic and motor territories of each participating brain region are functionally connected, explaining how abnormal activity in one region can affect activity at the network-level [55, 56]. Our findings show that striatal MSNs can reshape the functional connections within such a fronto - BG - cerebellar network and suggest a framework where D1 and D2 sub-populations differentially influence the network dynamics.

## Conclusion

Our study provides first evidence that the functional influence of striatal D1 and D2 MSNs on brain activity dynamics extends beyond the BG-thalamocortical network and is of differential rather than strictly antagonistic nature. Importantly, these effects seem to be dependent on their anatomical location in the striatum. In light of these findings, we propose that revised network models of BG function should take into consideration (i) cell-type specific influences (e.g. D1 vs D2 MSNs), (ii) the functional architecture of key brain areas within the BG (e.g. dm CPu vs vl CPu) and (iii) the involvement of regions beyond the BG-thalamocortical network (e.g. mPFC and cerebellum). Acknowledging these factors will pave the way for a more complete understanding of BG function within large-scale networks and have important implications for a wide range of movement and psychiatric disorders.

---

**Supplementary Figure 1.**
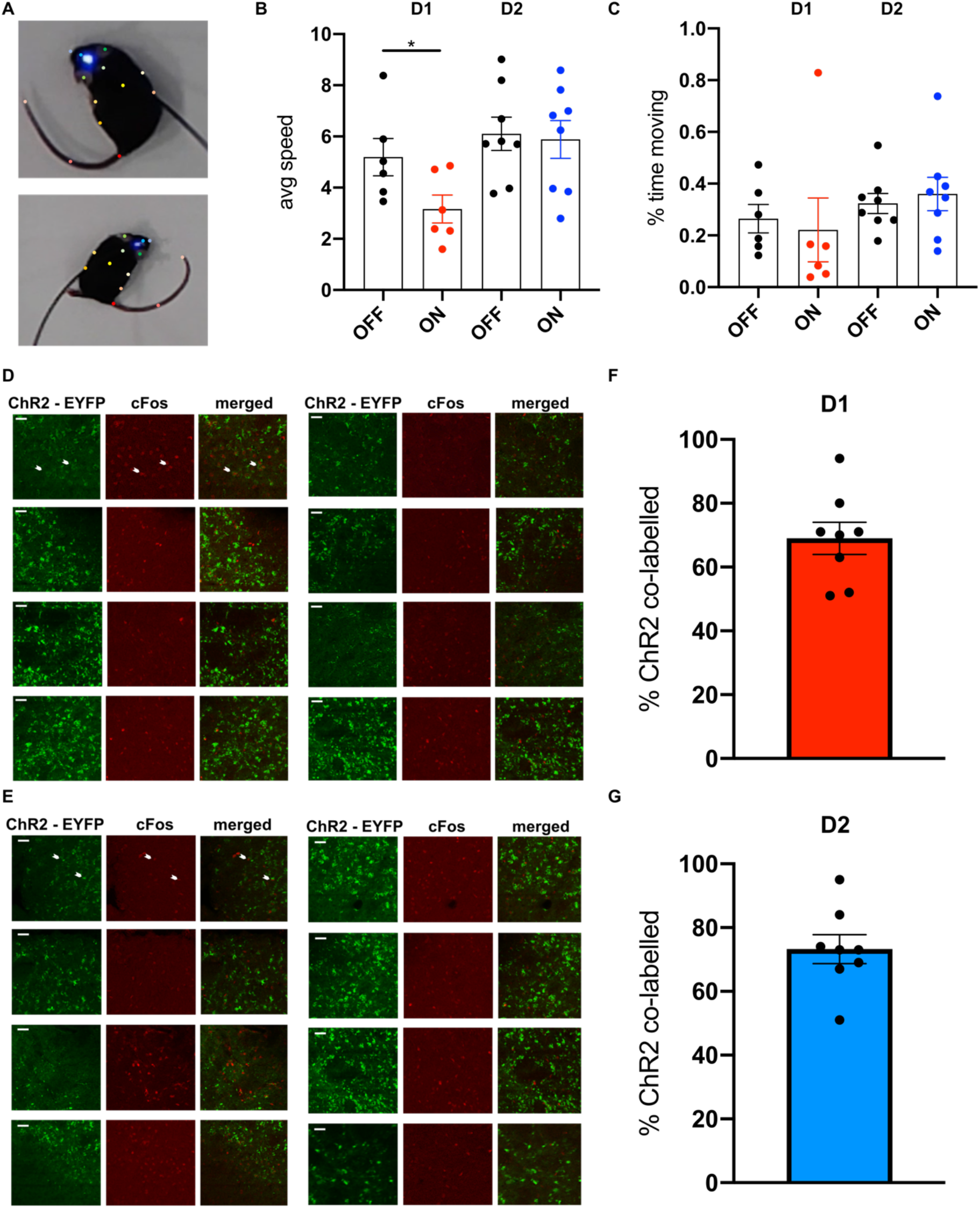
Optogenetic stimulation of striatal D1 and D2 MSNs elicits changes at the behavioural and molecular level. **A** Position of 13 body point labels for pose estimation using DeepLabCut. **B** Optogenetically evoked motor behaviour measured as changes in average speed. In D1-Cre mice, optogenetic MSN stimulation significantly decreases average speed (p = 0.046, linear mixed effects model with post hoc test), while no stimulation effect was detected in D2-Cre mice (p = 0.97, linear mixed effects model with post hoc test). **C** Optogenetically evoked motor behaviour measured as changes in percent of time moving. During optogenetic stimulation of striatal D1 and D2 MSNs no changes in the percent of time moving were detected (p(D1) = 0.43, p(D2) = 0.64, linear mixed effects model with post hoc test). **D** Histological verification of ChR2 and cFos expression 90 min after sustained laser stimulation in vl CPu D1 MSNs. White arrowheads indicate representative cells co-expressing ChR2-EYFP and the stained antibody cFos. **E** Histological verification of ChR2 and cFos expression 90 min after sustained laser stimulation in vl CPu D2 MSNs. White arrowheads indicate representative cells co-expressing ChR2-EYFP and the stained antibody cFos. **F** Quantification of cFos expression in ChR2-EYFP D1-Cre mice. **G** Quantification of cFos expression in ChR2-EYFP D2-Cre mice **p < 0.01, ***p < 0.001. Scale bars, 50μm.

**Supplementary Figure 2.**
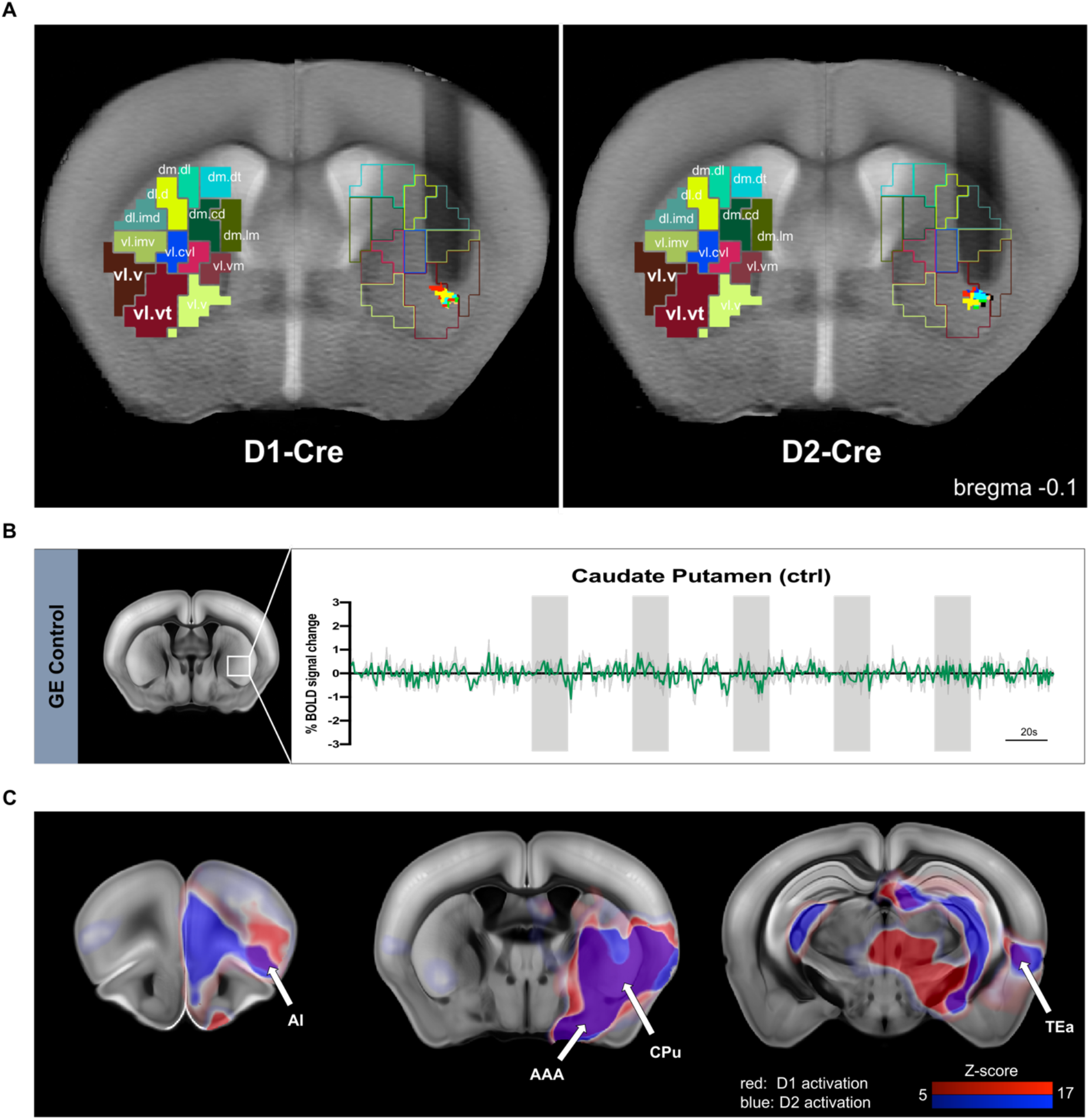
Optogenetic stimulation of D1 and D2 MSNs in the ventrolateral CPu drives brain-wide activation and de-activation hotspots. **A** Anatomical MRI verification of optical fiber placement in the vl CPu sub-region of D1-(n = 8 animals) and D2-Cre mice (n = 8 animals). **B** Mean time-series extracted from the right vl CPu of control animals depicted in green (n(D1) = 3 animals; no opsin expression). Laser stimulation blocks indicated in grey. **C** Overlay of thresholded D1 and D2 group z-stat activation maps, showing brain regions of similar BOLD signal profile. D1 group z-stat activation map displayed in red color code, D2 group z-stat activation map displayed in blue color code. Dm.dl, dorsomedial-dorsolateral; dm.d, dorsomedial-dorsal; dl.d, dorsolateral-dorsal; dl.imd, dorsolateral-intermediate; dm.cd, dorsomedial-central dorsal; dm.lm, dorsomedial-lateromedial; vl.imv, ventrolateral-intermedial ventral; vl.cvl, ventrolateral-central ventrolateral; vl.vm, ventrolateral-ventromedial; vl.v, ventrolateral-ventral; vl.vt, ventrolateral-ventral tip; AI, agranular insula; AAA, anterior amygdalar area; CPu, caudate putamen; TEa, temporal association area.

**Supplementary Figure 3.**
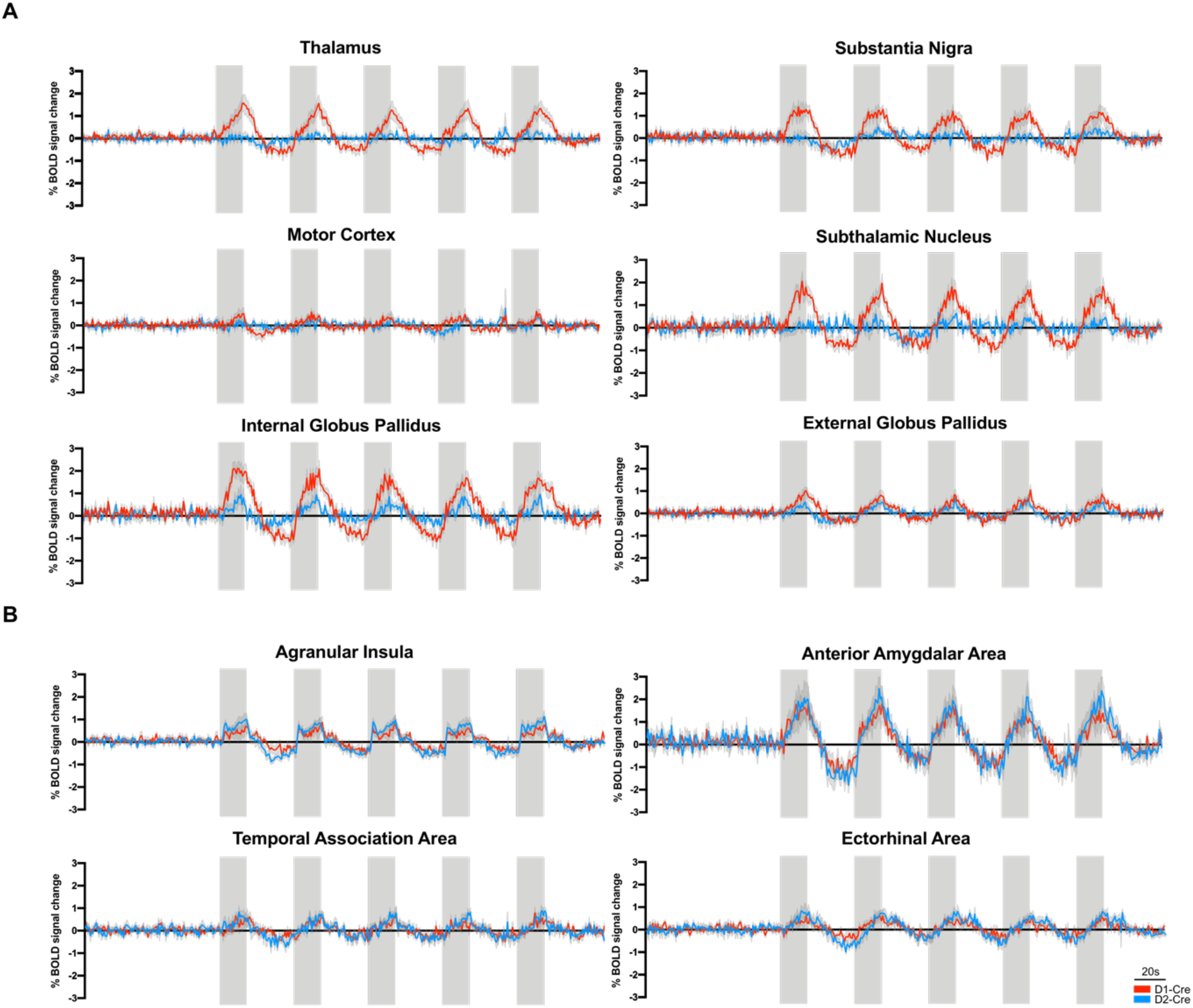
Mean time-series of GE BOLD local activation maxima and minima. **A** Mean time-series of D1-/D2-Cre mice extracted from selected BG regions of interest based on GE BOLD local activation maxima and minima z-stat maps. **B** Mean time-series of D1-/D2-Cre mice extracted from selected extra-BG regions of interest based on GE BOLD local activation maxima and minima z-stat maps. D1 MSN time-series depicted in red, D2 MSN time-series depicted in blue. Laser stimulation blocks indicated in grey.

**Supplementary Figure 4.**
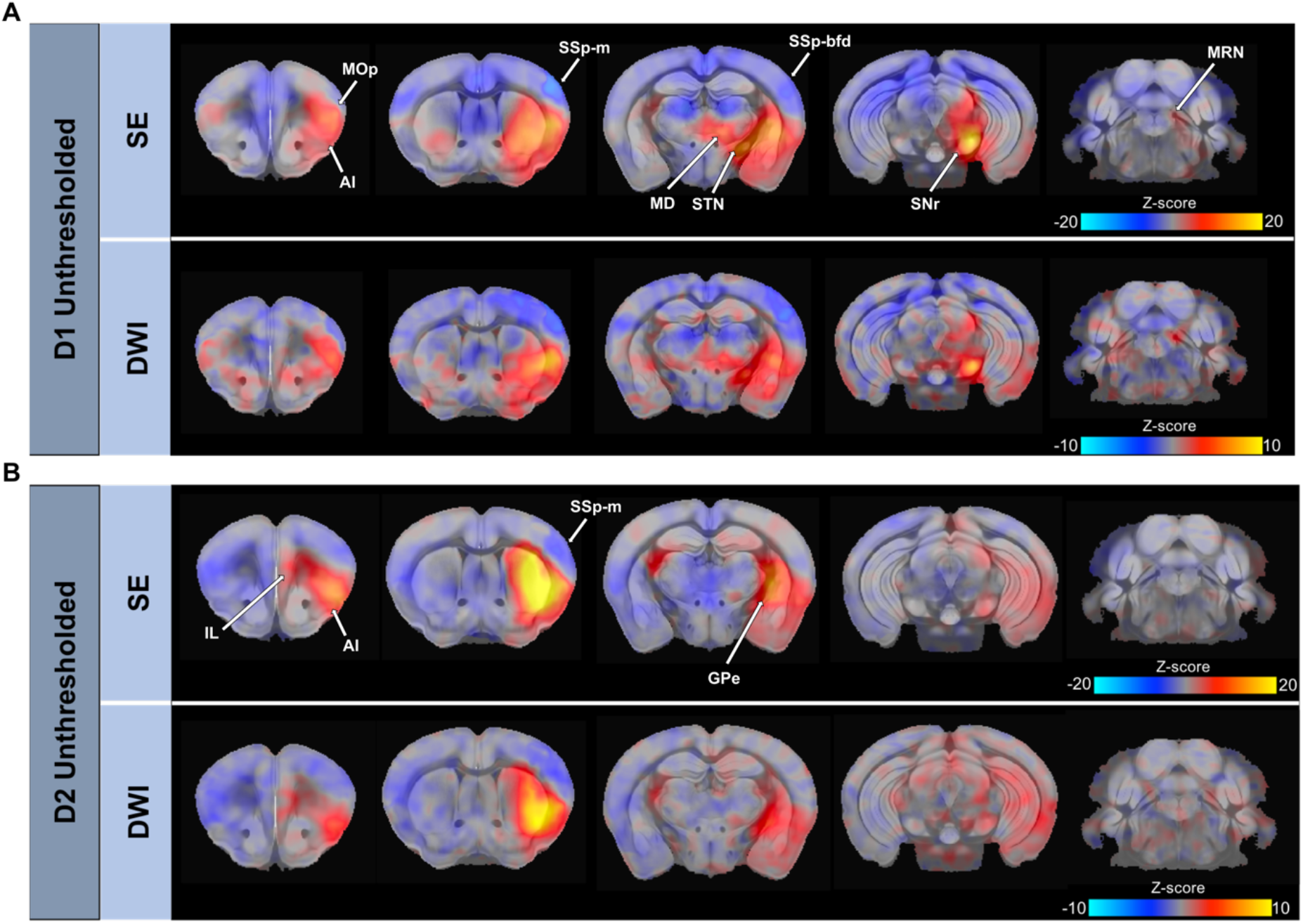
SE and dfMRI faithfully recapitulate causal influences of D1 and D2 MSN activity within the BG. **A** Unthresholded GLM spin-echo (SE) and dfMRI z-stat activation maps of D1 MSN stimulation. **B** Unthresholded GLM SE and dfMRI z-stat activation maps of D2 MSN stimulation. MOp, primary motor cortex; MD, mediodorsal nucleus of thalamus; STN, subthalamic nucleus; SSp-m, primary somatosensory cortex, mouth area; SSp-bfd, primary somatosensory cortex, barrel field; SNr, substantia nigra; MRN, midbrain reticular nucleus; IL, infralimbic area; AI, agranular insula; GPe, external globus pallidus.

**Supplementary Figure 5.**
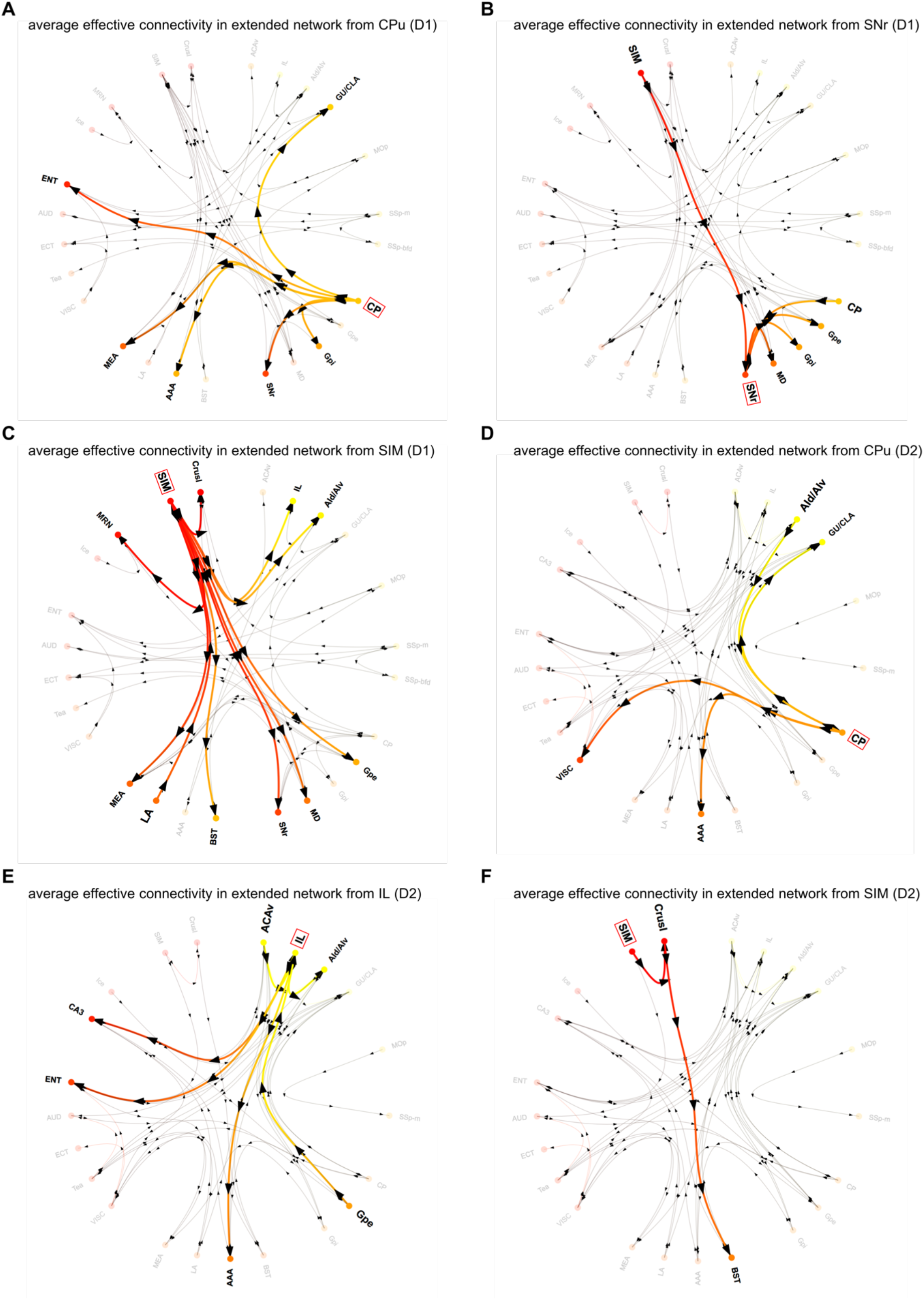
Whole-brain effective connectivity during D1 or D2 MSN stimulation as inferred using regression DCM (rDCM). **A** Average connectivity patterns in the extended network during D1 MSN stimulation, from the CPu, **B** from the SNr, and **C** from the SIM (Pp > 0.95). **D** Average connectivity patterns in the extended network during D2 MSN stimulation, from the IL, and **E** from the SIM (Pp > 0.95). Note, circos plots have been adjusted to show 50th percentile of increased directed connections only. AAA, anterior amygdalar area; ACAv, anterior cingulate; PrL, prelimbic area; AIv, ventral agranular insula; AUD, auditory area; BST, Bed nuclei of the stria terminalis; CA3, Ammon’s horn field 3; CPu, caudate putamen; ECT, ectorhinal area; ENT, entorhinal area; GPe, external globus pallidus; GPi, internal globus pallidus; GU/CLA, gustatory area/claustrum; Ice, inferior colliculus; IL, infralimbic area; LA, lateral amygdala nucleus; MEA, medial amygdala nucleus; MD, mediodorsal nucleus of thalamus; Mmme, medial part of the medial mammillary nucleus; MOp, primary motor cortex; MRN, midbrain reticular nucleus; SIM, simple lobule; SNr, substantia nigra pars reticulata; SBPV, subparaventricular zone; SSp-bfd, primary somatosensory cortex, barrel field; SSp-ll, primary somatosensory cortex, lower limb area; SSp-m, primary somatosensory cortex, mouth area; TEa, temporal association area; TU, tuberal nucleus; VISC, visceral area.

## Acknowledgments

We thank Jean-Charles Paterna from the Viral Vector Facility (VVF) of the Neuroscience Center Zürich, a joint competence center of ETH Zürich and the University of Zürich, for producing viral vectors and viral vector plasmids. C.G and V.Z are supported by the research grant ETH 062-18 and the Swiss National Science Foundation (SNSF) AMBIZIONE PZ00P3_173984/1. J.B. is supported by the ETH Zurich, ETH Project Grant ETH-20 19-1, SNSF Grant 310030_172889 and the 3R Competence Center. K.E.S. acknowledges support by the SNSF (project grant 320030_179377) and the René and Susanne Braginsky Foundation.

## Author Contributions

Conceptualization, C.G, S.F., J.B., K.E.S., D.R., N.W. and V.Z.; Methodology, C.G., S.F., C.S., L.v.Z, O.S., N.S. and V.Z.; Investigation, C.G., S.F., C.S., L.v.Z. and O.S.; Writing – Original Draft, C.G., S.F. and V.Z.; Writing – Review & Editing, C.G., S.F., C.S., L.v.Z., O.S., N.S., J.B., K.E.S., D.R., N.W. and V.Z.; Funding Acquisition, V.Z., J.B., K.E.S. and D.R.; Supervision, J.B., K.E.S., D.R., N.W. and V.Z.

